# The contribution of native protein complexes to targeted protein degradation

**DOI:** 10.1101/2025.06.17.660125

**Authors:** Lorraine Glennie, Nicole Curnutt, Tyrell Cartwright, Karen Dunbar, Brune Le Chatelier, Nicola T Wood, Thomas J Macartney, Christina M. Woo, Gopal P. Sapkota

## Abstract

Targeted protein degradation (TPD) destroys proteins of interest (POIs) by hijacking the cellular proteolytic machinery. Most proteins in cells exist and function as part of multi-protein or macromolecular complexes, thereby allowing a single protein to control multiple biological processes. Therefore, when a small molecule degrader induces proximity between an E3 ligase and the POI, the macromolecular context of the POI potentially influences the degradation outcomes of the POI and of the complex components. Here, we explore degradation of the eight CK1α-SACK1(A-H) (formerly known as FAM83A-H) complexes initiated by molecular glue degraders primarily designed to target Ser/Thr kinase CK1α. We demonstrate that lenalidomide-derived degraders DEG-77 and SJ3149, which selectively target the CK1α isoform, co-degrade multiple SACK1(A-H) proteins. We show that the degradation of SACK1(A-H) proteins by DEG-77 and SJ3149 requires CK1α, the CUL4A^CRBN^ E3 ligase complex and the proteasome. In cells derived from palmoplantar keratoderma patients harbouring the CK1α-binding deficient SACK1G^R265P^ mutation, DEG-77 targets CK1α and mitotic SACK1D but not SACK1G^R265P^, highlighting the requirement for CK1α-SACK1(A-H) interaction to achieve co-degradation. Our study underscores the importance of POI context in TPD and reinforces the potential for selectively targeting specific protein complexes for degradation.

## Introduction

A protein’s function within a cell is shaped by its subcellular localization, interacting partners, post-translational modifications and turnover dynamics ^1,2^. As such, protein-protein interactions and multi-protein complexes constitute the fundamental functional modules that orchestrate cellular signalling and homeostasis ^3,4^. A single protein can participate in diverse cellular processes by forming unique macromolecular complexes depending on the context. The Ser/Thr protein kinase CK1α, a prototypic member of the CK1 kinase family, exemplifies this principle. CK1α regulates numerous cellular processes including WNT signalling, cell division, apoptosis, calcium signalling, and cellular responses to DNA damage ^5–9^. While the mechanisms underlying its broad activity are still being elucidated, the eight proteins of the FAM83 family (A-H; hereafter referred to as SACK1A-H) are recognized as key regulators of CK1α and other CK1 isoforms ^10–13^. They interact with CK1α through the conserved Scaffold Anchor of CK1 (SACK1) domain (formerly known as DUF1669), dictating its subcellular localization and potentially its substrate repertoire ^11,13^. SACK1D recruits CK1α to the mitotic spindle to orchestrate proper spindle alignment and timely cell division ^6^. The CK1α-SACK1F complex at the plasma membrane and CK1α-SACK1G complex primarily in the cytoplasm drive the activation of canonical WNT signalling ^10,12^. Indeed, loss of CK1α-SACK1G interaction, leading to inhibition of canonical WNT signalling, underlies the pathogenesis of palmoplantar keratoderma (PPK) caused by autosomal recessive *FAM83G* mutations in both humans ^14–16^ and dogs ^17^. Four reported missense *FAM83G* mutations in PPK, which lead to A34E, R52P, F58I & R265P substitutions on the SACK1G protein, all lie within the SACK1 domain and disrupt CK1α binding ^14–16,18^.

Targeted protein degradation (TPD), via proteolysis-targeting chimeras (PROTACs), molecular glue degraders (MGDs), or other modalities, is emerging as a powerful approach for investigating protein function in cells and developing targeted therapeutics ^19–23^. PROTACs and MGDs function by inducing spatial proximity between an E3 ubiquitin (Ub) ligase and the target protein of interest (POI), thereby facilitating POI ubiquitylation and subsequent proteasomal degradation ^24–26^. Excitingly, several PROTACs have progressed to human clinical trials, whereas some MGDs based on immunomodulatory imide drugs (IMiDs), such as thalidomide, lenalidomide and pomalidomide, are already in clinical use against multiple myeloma and other indications ^27–29^. IMiDs act as MGDs by binding to the CUL4A E3 ligase substrate receptor CRBN and facilitate the recruitment of neo-substrates, which then undergo ubiquitylation and proteasomal degradation ^24,25,30,31^. Naturally, CRBN is thought to specifically recognize proteins with a C-terminal cyclic imide degron, which arises from the cyclization of glutamine or asparagine residues ^32^. Many neo-substrates recruited to CRBN by IMiD-based MGDs contain C2H2 zinc finger domains, such as transcription factors IKZF1 and IKZF3 ^25,33,34^. Proteins like CK1α are also recruited by lenalidomide as a neo-substrate through its exposed β-hairpin motif in the N-lobe of the kinase domain ^24,35^. Degradation of CK1α is thought to contribute to the clinical efficacy of lenalidomide against myelodysplastic syndromes (MDS) caused by the deletion of chromosome 5q that results in the loss of one *CSNK1A1* (CK1α) allele ^24^. Successful clinical applications of IMiDs have rejuvenated profound interest in developing even more potent and selective CRBN ligands that target C2H2 zinc-finger containing proteins as well as those that contain exposed β-hairpin motifs similar to that of CK1α ^33,34,36^.

Independent efforts to improve selectivity and potency of CK1α degradation with lenalidomide-derivatives have led to novel series of potent CK1α degraders. The DEG series of compounds derived from lenalidomide resulted in DEG-77 and DEG-35 as potent dual CK1α and IKZF2 degraders with efficacy against acute myeloid leukaemia ^37,38^. DEG-77 also displayed antiproliferative activity against diffuse large B cell lymphoma cell line OCI-LY3 and the ovarian cancer cell line A2780 ^38^. Independently developed lenalidomide-derivatives SJ7095, SJ0040 and SJ3149 are also potent CK1α degraders ^39,40^. While SJ7095 degraded CK1α, IKZF1 and IKZF3, and caused cytotoxicity against acute myeloid leukaemia MOLM-13 cells, SJ0040 and SJ3149 improved selectivity against CK1α but still maintained antiproliferative effects ^40^.

TPD in cells is primarily driven by the generation of a productive POI-degrader-E3 ternary complex. Beyond chemical properties of the degrader, there are many cellular factors that can influence the formation of this ternary complex and thus the specificity and potency of the degrader ^41–43^. For example, subcellular localization of the POI and/or E3 ligase, the choice of the E3 ligase being recruited and the abundance and activity of the E3 ligase can all influence the extent and kinetics of TPD ^42,44,45^. Other cellular factors such as degrader uptake and retention, degrader metabolism and protein turnover rates can also affect TPD ^41^. One crucial, and often overlooked, factor in TPD that can influence the formation of POI-degrader-E3 ternary complex is the inherent POI macromolecular context that defines its interactions with other macromolecules, localization and activity. Based on this POI context, the extent and kinetics of POI degradation as well as potential degradation of other proteins in complex with the POI, often referred to as *‘bystander degradation’* or *‘collateral damage’*, could be affected.

Previously, we showed that lenalidomide, which only modestly degrades CK1α in cells, leads to complete co-degradation of SACK1F but not other SACK1 proteins ^31^. Given recent reports of potent and selective CK1α degraders ^37–40^ and our findings that CK1α in cells exists within a unique subcellular location dictated by its interaction with one of eight SACK1A-H proteins ^11^, we postulated that these degraders could also influence specific or multiple CK1α-SACK1A-H complexes. Here, we explore the effects of DEG ^38^ and SJ ^40^ series of potent CK1α degraders on the stability of SACK1A-H proteins.

## Results

### Assessment of endogenous CK1α:SACK1(A-H) complexes in multiple human cell lines

The eight SACK1A-H proteins are known to direct the localization of the Ser/Thr kinase CK1α to distinct subcellular compartments (**Fig. 1A**) ^11,13^. We monitored the expression of CK1α and different SACK1A-H proteins and their interactions at the endogenous level in several human cancer cell lines. In DLD1 colorectal cancer cells, the expression of CK1α, SACK1B, SACK1D, SACK1F, SACK1G and SACK1H was detected by immunoblotting (**Fig. 1B**). In U2OS osteosarcoma cells, CK1α, SACK1D, SACK1G and SACK1H were detected, while in A549 lung adenocarcinoma cells only CK1α, SACK1G and SACK1H were detected (**Fig. 1B**). In TOV-21G ovarian cancer cells, CK1α, SACK1B, SACK1D, SACK1G and SACK1H were detected (**Fig. 1C**). We currently lack antibodies to detect SACK1A, SACK1C and SACK1E at the endogenous level. Immunoprecipitates (IPs) of endogenous CK1α with anti-CK1α antibody co-precipitated SACK1B, SACK1F, SACK1G and SACK1H from DLD1 extracts, SACK1G and SACK1H from U2OS and A549 extracts, and SACK1B, SACK1G and SACK1H from TOV-21G extracts (Fig. 1B). No SACK1D was co-precipitated in anti-CK1α IPs from any of the extracts which were harvested under asynchronous conditions (**Fig. 1B&C)**. This is consistent with our previous findings that SACK1D-CK1α interaction occurs only during mitosis and not in other stages of the cell cycle ^6^. As expected, IPs with IgG controls did not precipitate CK1α or any SACK1 protein (**Fig. 1B&C)**. Using these cells, we sought to test the impact of CK1α degraders on different SACK1 proteins.

**Figure 1:**
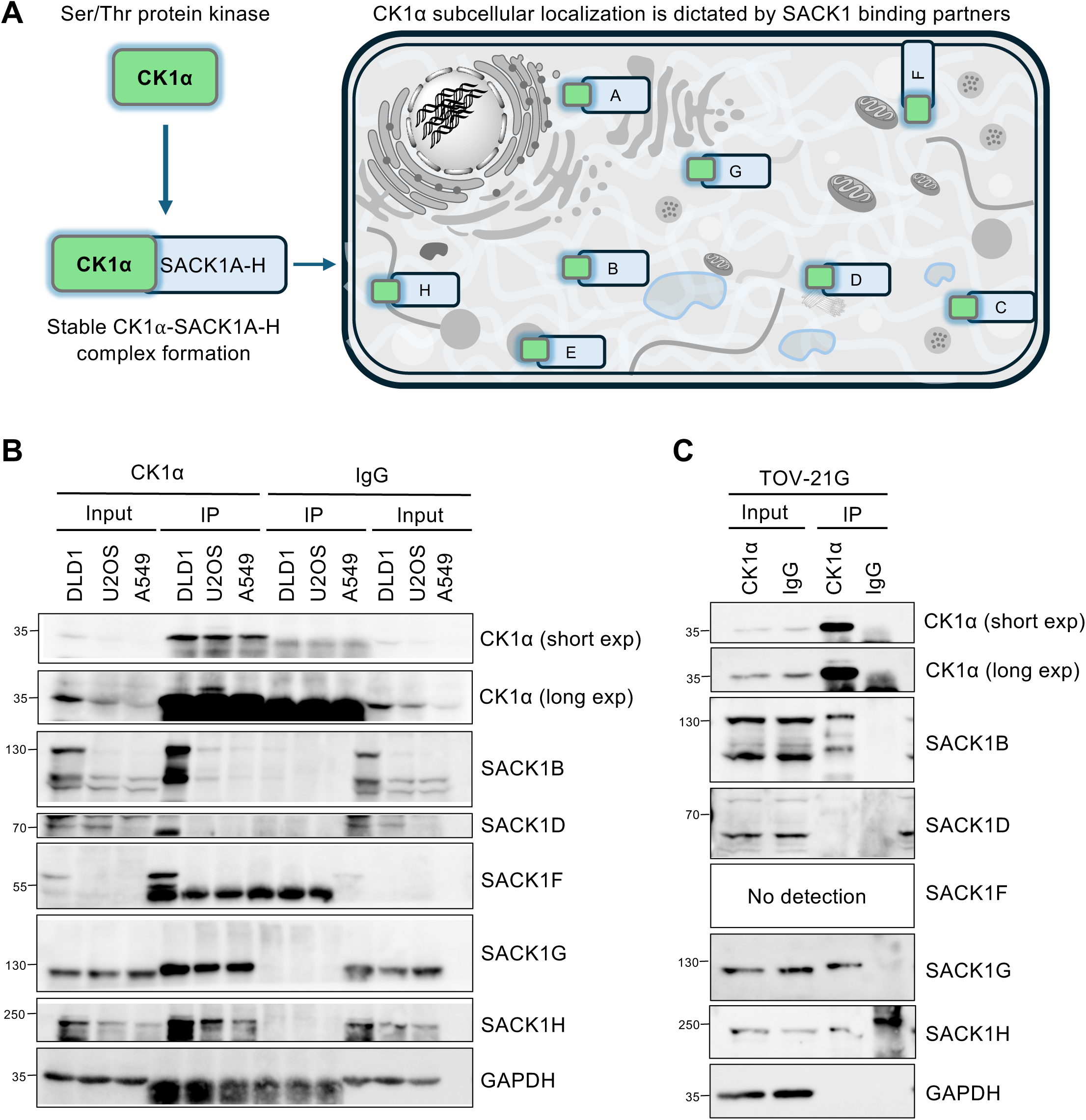
SACK1 domain-containing proteins form complexes with CK1α at the endogenous level in DLD1, U2OS, A549 and TOV-21G cell lines. **(A)** Depiction of subcellular distribution of serine/threonine protein kinase CK1α and interacting SACK1 domain-containing proteins. SACK1(A-H) direct CK1α and other CK1 isoforms to distinct subcellular compartments to regulate CK1 biology. **(B&C)** DLD1, U2OS, A549 (B) and TOV-21G (C) cells were lysed, and extracts normalised to 2 mg total protein and subjected to anti-CK1α IPs. IgG IPs were used as negative controls. Input extracts and IPs were resolved by SDS-PAGE, transferred to PVDF membrane and analysed by immunoblotting using the indicated antibodies.

### DEG series of CK1α degraders reduce levels of several SACK1 proteins

Given that we detected CK1α and most SACK1(A-H) proteins in DLD1 cells, we treated these cells with the DEG series ^37,38^ of CK1α degraders **(Fig. S1)**. DEG compounds (DEG-35, −48, −52, −54, −61, −64 and −77) **(Fig. S1)** were selected due to their range of efficacy of CK1α degradation (low nanomolar to micromolar DC_50, CK1α_) and structural diversity within the DEG series, which we reasoned may lead to differences in destabilization of specific CK1α-SACK1(A-H) complexes upon compound treatment. These compounds were added to cells at a concentration of 100 nM for 24 h (**Fig. 2A**). CK1α, SACK1F and SACK1G were robustly depleted by DEG-35 and DEG-77 with a >90% reduction in observed protein abundance compared to DMSO-treated control (**Fig. 2A**). DEG-52 and DEG-64 led to modest CK1α and SACK1G depletion but almost complete removal of SACK1F (**Fig. 2A**). DEG-54, DEG-48 and DEG-61 led to only a slight decrease in the level of CK1α and SACK1G but a robust decrease in SACK1F (**Fig. 2A**). SACK1D interacts with CK1α only in mitosis, and this results in its hyperphosphorylation, causing a mobility shift of ∼25 kDa in immunoblot assays ^11^. In DMSO-treated asynchronous DLD1 extracts, we detected two differentially migrating SACK1D signals, an intense lower (∼75 kDa) band and a less intense upper (∼100 kDa) band, suggesting a proportion of mitotic cells in our experimental conditions. The lower band represents unphosphorylated SACK1D protein that does not interact with CK1α, whereas the upper band represents the mitotic hyperphosphorylated SACK1D bound to CK1α ^6^. DEG-35, DEG-54, DEG-52, DEG-61 and DEG-77 led to a substantial reduction in levels of the 100 kDa hyperphosphorylated SACK1D without affecting the levels of the 75 kDa unphosphorylated SACK1D, whereas DEG-64 resulted in reduction in levels of both pools (**Fig. 2A**). There was only a modest reduction in levels of SACK1B by DEG-35, DEG-54, DEG-52, DEG-64 and DEG-77 but no changes by other DEG compounds (**Fig. 2A**). SACK1H levels did not change substantially by any of the DEG compounds (**Fig. 2A**). While SACK1D, SACK1F and SACK1G bind selectively to CK1α, SACK1B and SACK1H also bind to CK1δ and CK1ε, which are not targeted for degradation by imide compounds.

**Figure 2:**
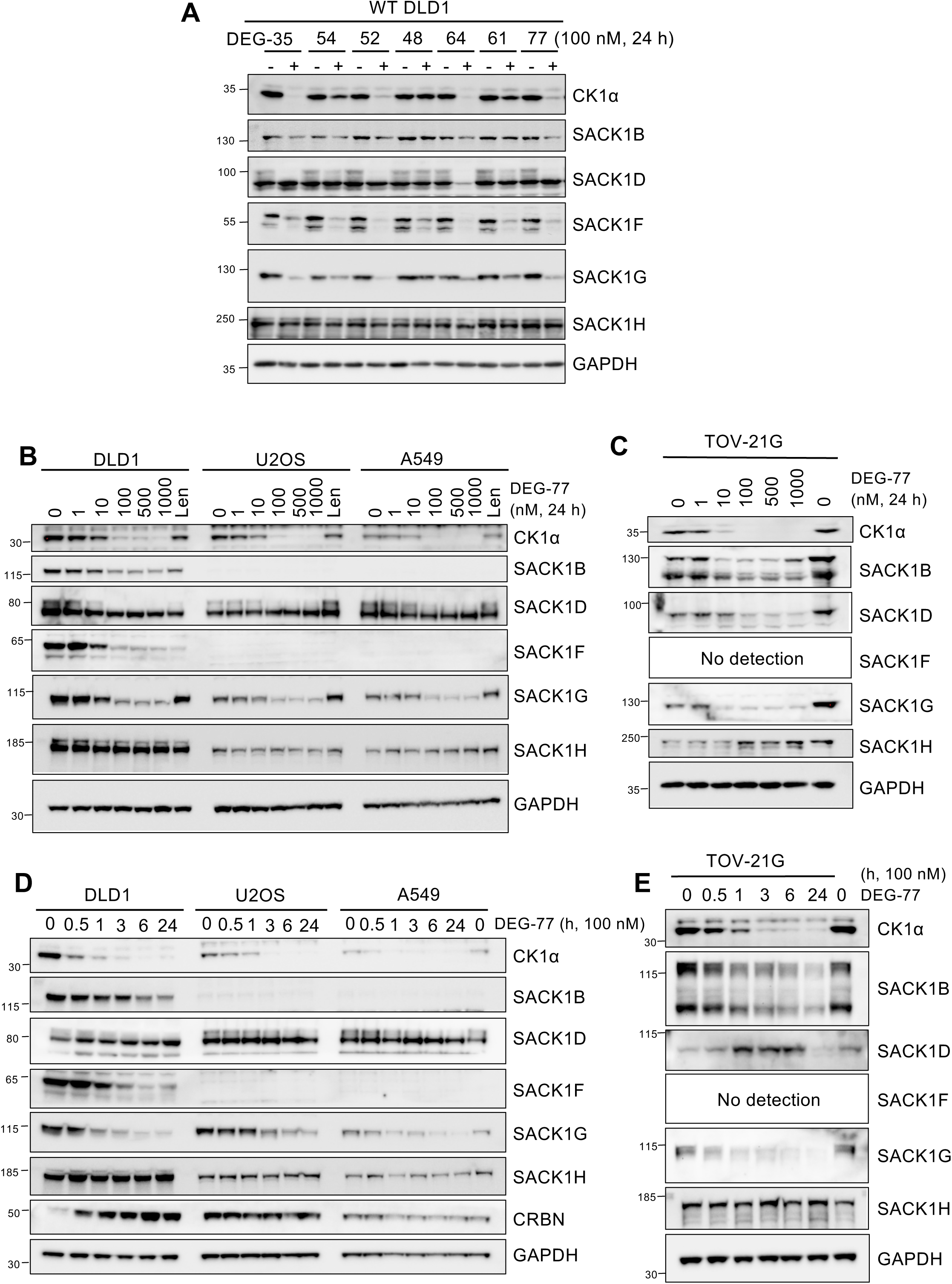
DEG-series of CK1α degraders lead to co-depletion of CK1α-SACK1(A-H) complexes to varying degrees. **(A)** Wild-type DLD1 cells were incubated with DMSO or 100 nM of each DEG compound (−35, −54, −52, −48, −64, −61, −77) as indicated for 24 h before cell lysis. Extracts were resolved by SDS-PAGE, transferred to PVDF membrane and immunoblots were incubated with the indicated antibodies. **(B)** Wild-type DLD1, U2OS and A549 cell lines were incubated with increasing concentrations of DEG-77 (1-1000 nM) or 10 µM lenalidomide for 24 h before cell lysis. 20 µg extract protein was resolved by SDS-PAGE and transferred to PVDF membranes for immunoblotting analysis using the indicated antibodies. **(C)** As in (B), except TOV-21G cells were cultured for 24 h with increasing concentrations of DEG-77 (1-1000 nM) before cell lysis. **(D)** As in (B), except wild-type DLD1, U2OS, and A549 cells were incubated with 100 nM DEG-77 for different time periods (0.5 - 24 h) before cell lysis. **(E)** As in (C), except TOV-21G cells were incubated with 100 nM DEG-77 for different time periods (0.5 - 24 h) before cell lysis. All blots are representative of at least 3 biological replicates.

Next, to determine if DEG-77 co-depletes CK1α and interacting SACK1B, SACK1D, SACK1F, SACK1G and SACK1H proteins concurrently in multiple cells, DLD1, U2OS, A549 and TOV-21G cells were treated with an increasing concentration of DEG-77 (1-1000 nM) for 24 h (**Fig. 2B&C)**. The parental compound lenalidomide (10 μM), from which DEG-77 was derived, was also employed as a control. DEG-77 induced a dose-dependent co-depletion of CK1α, SACK1B, hyperphosphorylated SACK1D, SACK1F and SACK1G but not SACK1H compared to DMSO control in all cells that expressed these proteins, with a robust reduction observed at concentrations of 100 nM or over (**Fig. 2B&C)**. Lenalidomide caused a minimal degradation of CK1α and no decrease in levels of SACK1B, SACK1D, SACK1G and SACK1H but a robust decrease in SACK1F levels compared to DMSO control (**Fig. 2B**). When cells were treated with 100 nM DEG-77 over a course of 24 h, a time-dependent reduction in levels of CK1α, SACK1B, hyperphosphorylated SACK1D, SACK1F and SACK1G but not SACK1H was observed (**Fig. 2D&E)**. A reduction in levels of CK1α was observed as early as 30 min following DEG-77 treatment but a robust depletion was observed after 1 h, with peak degradation observed at 3 h, which was sustained for 24 h (**Fig. 2D&E)**. The decreased level of SACK1B in DLD1 and TOV-21G cells, hyperphosphorylated SACK1D in DLD1, U2OS and A549 cells, SACK1F in DLD1 cells and SACK1G in all cells closely mirrored the degradation kinetics of CK1α (**Fig. 2D&E)**.

### SJ series of CK1α degraders reduce levels of several SACK1 proteins

Similar to the DEG series of compounds above, we next investigated the SJ series of CK1α degraders ^39,40^ **(Fig. S1)** for their ability to co-deplete SACK1(A-H) proteins. We treated DLD1 cells with increasing concentrations from 1-1000 nM of SJ7095, SJ0040 and SJ3149 for 4 h (**Fig. 3A**). All compounds led to a dose-dependent decrease in levels of CK1α, SACK1B, SACK1F and SACK1G, with reduction observed at doses over 100 nM (**Fig. 3A**). SJ7095 was the least potent CK1α degrader, with reduced CK1α levels observed at 100 nM and optimal degradation occurring at 1-10 μM (**Fig. 3A**). SJ0040 and SJ3149 were both potent CK1α degraders, with modest degradation observed at 10 nM and almost complete CK1α degradation observed at concentrations over 100 nM (**Fig. 3A**). Despite the differences in the extent of CK1α degradation by these compounds, there was no substantial difference observed in the reduction of SACK1B, SACK1F and SACK1G (**Fig. 3A**). The improved potency of CK1α degradation by SJ0040 and SJ3149 in comparison to SJ7095 is potentially due to their stabilised interaction with CK1α within the CK1α-imide-CRBN complex ^40^.

**Figure 3:**
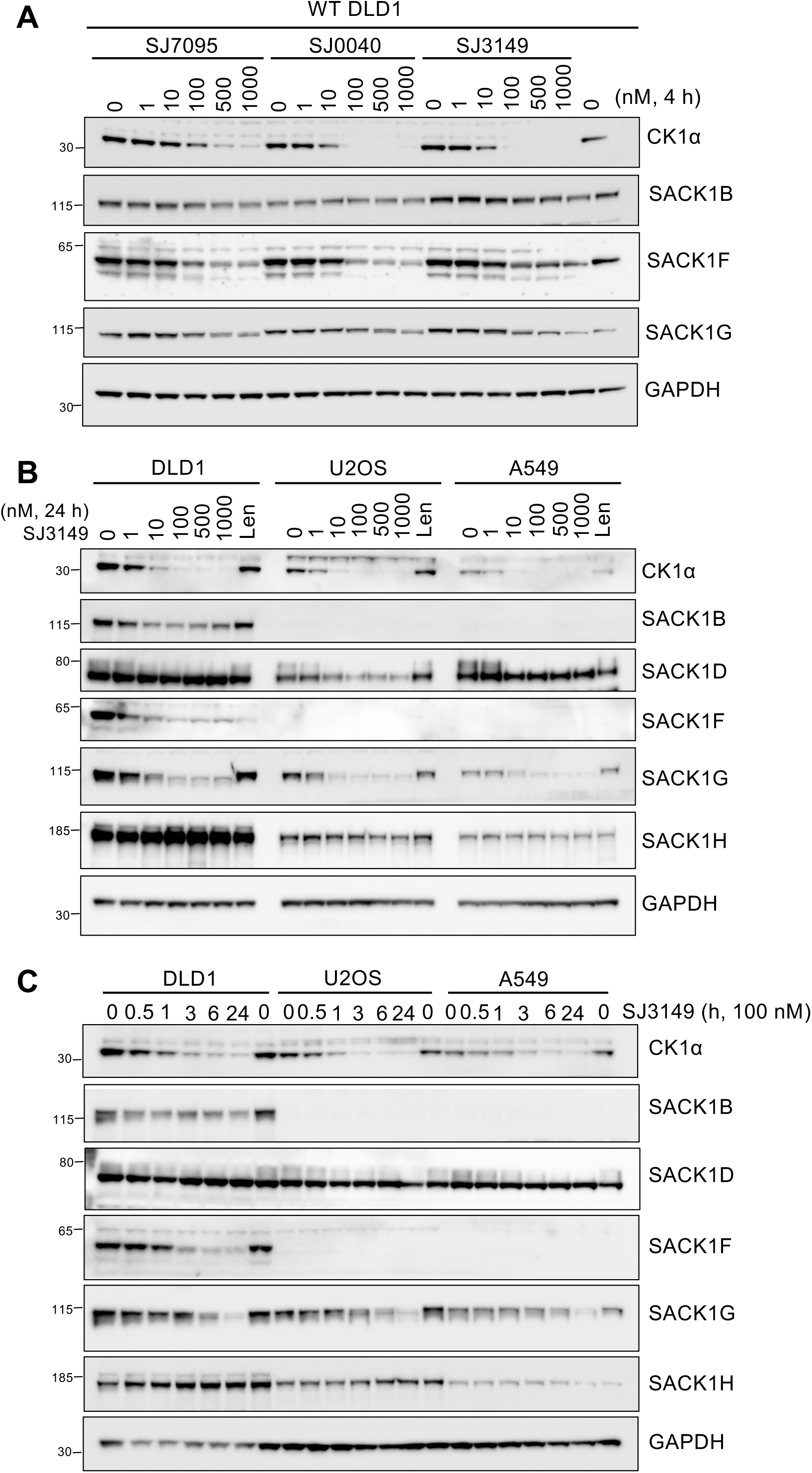
SJ-series of CK1α degraders lead to co-depletion of CK1α-SACK1(A-H) complexes. **(A)** Wild-type DLD1 cells were treated with SJ compounds (SJ7095, SJ0040, SJ3149) at increasing concentrations from 1-1000 nM for 4 h before cell lysis. **(B)** Wild-type DLD1, U2OS and A549 cell lines were incubated with increasing concentrations of SJ3149 (1-1000 nM) or 10 µM lenalidomide for 24 hours before cell lysis. 20 µg extract protein was resolved by SDS-PAGE and transferred to PVDF membranes for immunoblotting analysis using the indicated antibodies. **(C)** As in (B) except wild-type DLD1, U2OS, and A549 cells were incubated with 100 nM SJ3149 for different time periods (0.5 - 24 h) as indicated before cell lysis. All blots are representative of at least 3 biological replicates.

Next, we treated DLD1, U2OS and A549 cells with increasing doses (1-1000 nM) of SJ3149 for 24 h (**Fig. 3B**). Similar patterns of co-depletion in levels of CK1α, SACK1B (in DLD1 cells), hyperphosphorylated SACK1D, SACK1F (in DLD1 cells), and SACK1G was observed compared to DMSO control (**Fig. 3B**). Some depletion was observed with 10 nM SJ3149, with maximal depletion observed at higher (100-1000 nM) concentrations (**Fig. 3B**). No substantial depletion in levels of SACK1H was observed in any cells (**Fig. 3B**). A time-course treatment of 100 nM SJ3149 over 24 h revealed a time-dependent co-depletion in levels of CK1α, SACK1B (in DLD1 cells), hyperphosphorylated SACK1D, SACK1F (in DLD1 cells) and SACK1G, with some depletion observed as early as 30 min following SJ3149, and maximal depletion achieved after 3-6 h, which was sustained until 24 h (**Fig. 3C**). The kinetics and potency of co-degradation of CK1α and the interacting SACK1 proteins by SJ3149 and DEG-77 were similar (**Fig. 2&3)**.

### Comparison of CK1α degraders for their ability to co-deplete SACK1 proteins

Given the differences in co-depletion of CK1α and different interacting SACK1 proteins between lenalidomide, DEG-48, DEG-77 and SJ3149, we assessed these CK1α degraders together with a previously reported CK1α and CDK7/9 degrader BTX161 ^46^ to compare how specific CK1α-SACK1(A-H) complexes were co-depleted by each. DLD1 cells were treated with lenalidomide (10 µM), BTX161 (10 µM), DEG-48 (100 nM), DEG-77 (100 nM), SJ7095 (1 µM) and SJ3149 (100 nM) or DMSO control for 24 h (**Fig. 4A&B)**. BTX161, DEG-77 and SJ3149 caused a robust degradation of CK1α, while lenalidomide and SJ7095 caused moderate degradation and DEG-48 did not cause any CK1α degradation (**Fig. 4A&B)**. For potent degraders, DEG-77 and SJ3149 caused robust co-depletion of SACK1B, SACK1F and SACK1G (**Fig. 4A&B)**. BTX161 and lenalidomide only robustly co-depleted SACK1F but no other SACK1 protein (**Fig. 4A&B)**. SJ7095 caused a robust depletion of SACK1F but only a moderate depletion of SACK1G and no degradation of SACK1B and SACK1H (**Fig. 4A&B)**. These findings suggest that despite all IMiD compounds engaging CRBN, the specific nature by which each degrader recruits specific CK1α-SACK1(A-H) complex to the CUL4A^CRBN^ E3 ligase machinery potentially dictates how efficiently each complex can be targeted for degradation.

**Figure 4:**
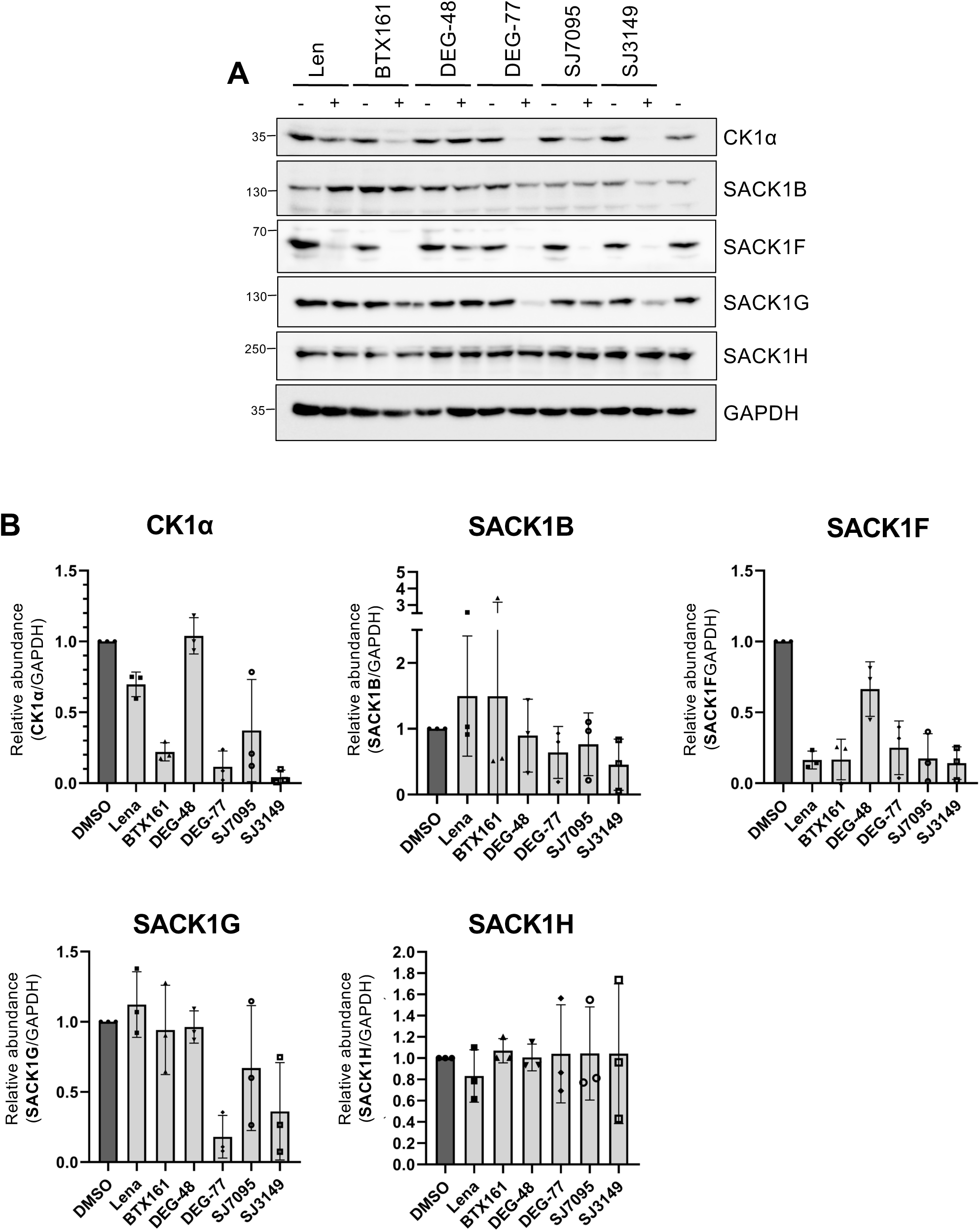
Comparison of CK1α degraders for their ability to co-deplete the interacting SACK1 proteins. **(A)** Wild-type DLD1 cells were incubated with different imide-derived CK1α degraders: lenalidomide (10 µM), BTX161 (10 µM), DEG-48 (100 nM), DEG-77 (100 nM), SJ7095 (1 µM) and SJ3149 (100 nM) or DMSO control for 24 h before cell lysis. 20 µg extract protein was resolved by SDS-PAGE, transferred to PVDF membrane and analysed by immunoblotting using the indicated antibodies. Blots are representative of at least 3 biological replicates. **(B)** Quantification by densitometry of protein signals shown in (A) using Fiji 1.53q (ImageJ). Each graph depicts the abundance of CK1α or SACK1(A-H) proteins normalised to GAPDH loading control relative to DMSO treatment control (n = 3, error bars represent mean ± SD).

### The degradation of SACK1 proteins by DEG-77 and SJ3149 requires CK1α, CRBN and the proteasome

To examine if the observed co-depletion of CK1α-SACK1(A-H) complexes by imides is mediated through the recruitment of CK1α to CRBN (**Fig. 5A**), we generated CSNK1A1^-/-^ **(Fig. S2)** and CRBN^-/-^ ^31^ DLD1 cells by CRISPR/Cas9 genome editing (**Fig. 5B&C)**. Wild-type, CRBN^-/-^ and CSNK1A1^-/-^ DLD1 cells were treated with 100 nM DEG-77 (**Fig. 5B**) or SJ3149 (**Fig. 5C**) for 6 h. By immunoblotting, no CK1α protein was detected in CSNK1A1^-/-^ cells compared to both wild-type and CRBN^-/-^ DLD1 cells. Similarly, in comparison to levels detected in WT and CSNK1A1^-/-^ DLD1 cells, CRBN levels were almost absent in CRBN^-/-^ DLD1 cells, although a slight residual signal remained (**Fig. 5B&C)**. As expected, the degradation of CK1α by DEG-77 (**Fig. 5B**) and SJ3149 (**Fig. 5C**) observed in WT DLD1 cells was rescued in CRBN^-/-^ DLD1 cells. Excitingly, the depletion in levels of SACK1B, SACK1F and SACK1G caused by DEG-77 (**Fig. 5B**) and SJ3149 (**Fig. 5C**) observed in WT DLD1 cells was completely rescued in both CRBN^-/-^ and CSNK1A1^-/-^ DLD1 cells, suggesting that the imide-induced depletion in levels of SACK1 proteins requires both CRBN and CK1α. We observed decreased levels of SACK1G in CSNK1A1^-/-^ DLD1 cells compared to WT or CRBN^-/-^ cells (**Fig. 5B&C)**. This is consistent with our previous observations where interaction between SACK1G and CK1α is essential for regulating the stability of each protein and depleting one or disrupting their interaction destabilizes the other ^12,14,16^. To determine how CK1α-SACK1(A-H) co-degradation was mediated by DEG-77 and SJ3149, wild-type DLD1 cells were pre-treated with either the proteasomal inhibitor MG132 (20 µM) or autophagy-lysosomal inhibitor bafilomycin A1 (50 nM) for 2 h prior to treatment with DEG-77 and SJ3149 (**Fig. 5D&E)**. The depletion in levels of CK1α, SACK1B, SACK1F or SACK1G by DEG-77 and SJ3149 was completely rescued when cells were pre-treated with MG132 but not bafilomycin A1 (**Fig. 5D&E)**, indicating that CK1α-SACK1 complex co-degradation by DEG-77 and SJ3149 treatment is mediated by the proteasome.

**Figure 5:**
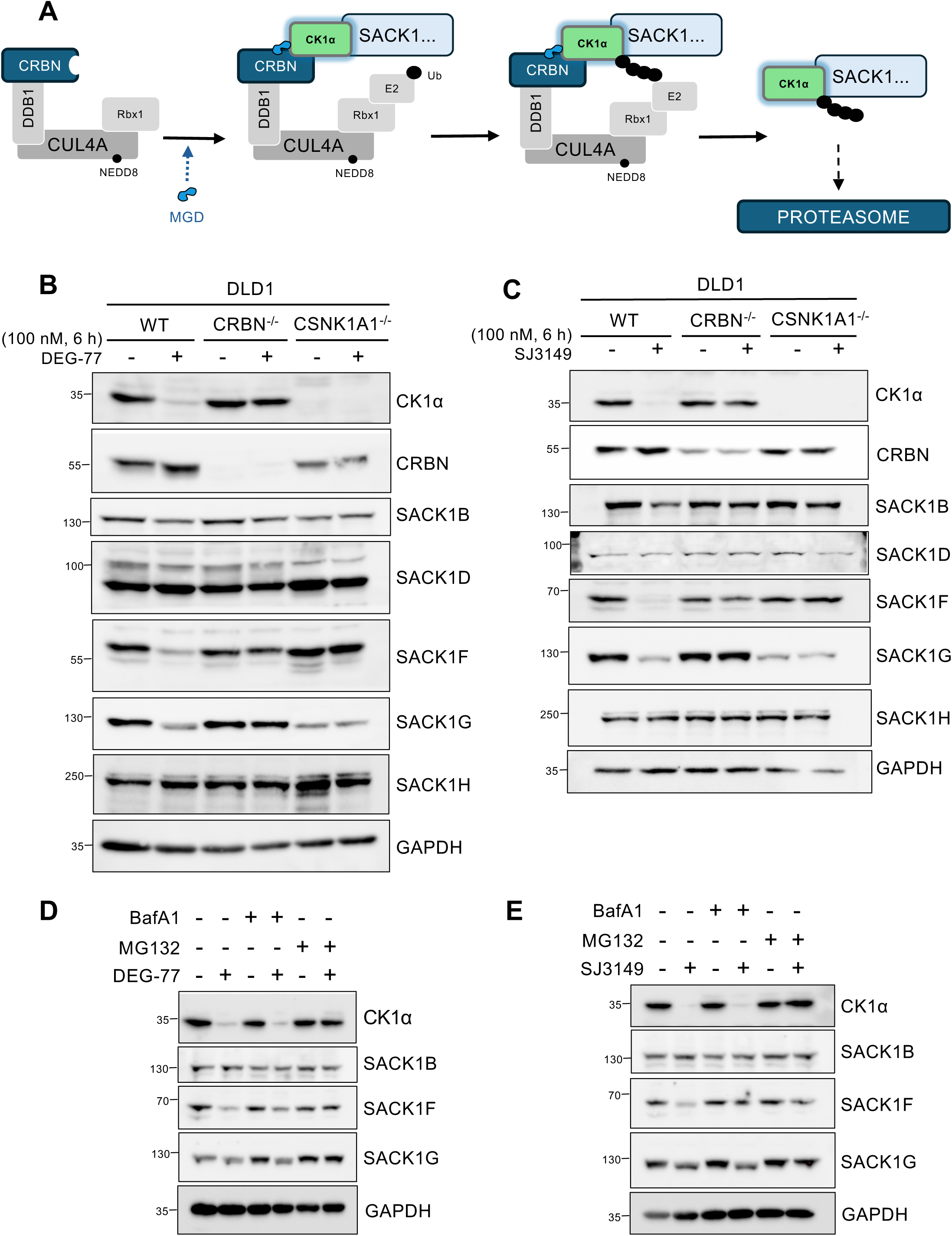
The degradation of SACK1 proteins by imides DEG-77 and SJ3149 requires CK1α, CRBN and the proteasome. **(A)** The postulated mechanism of action by which imide-derived molecular glue degraders (MGDs) DEG-77 and SJ3149 induce CK1α-SACK1(A-H) co-degradation. By binding to substrate receptor cereblon (CRBN), both MGDs primarily recruit CK1α to CUL4^CRBN^ E3 ligases and, as CK1α exists in complex with SACK1(A-H) proteins, it co-recruits SACK1(A-H) proteins to the CUL4^CRBN^ E3 ligase complex. Depending on the proximity of the recruited CK1α-SACK1(A-H) complex to the CUL4^CRBN^ E3 catalytic site, specific CK1α-SACK1(A-H) complexes are potentially ubiquitylated and targeted for degradation via the proteasome. **(B&C)** Wild-type, CRBN^-/-^ and CSNK1A1^-/-^ DLD1 cells were incubated with 100 nM DEG-77 or DMSO (B) or 100 nM SJ3149 or DMSO (C) for 6 h before cell lysis. 20 µg extract protein was resolved by SDS-PAGE and transferred to PVDF membrane for immunoblotting analysis. **(D&E)** As in (B), except wild-type DLD1 cells were pre-treated with either MG132 (20 µM) or bafilomycin A1 (50 nM) for 2 hours before treatment with DEG-77 (100 nM) or DMSO (D) or SJ3149 (100 nM) or DMSO (E) for a further 4 h before lysis. For (B-E), blots are representative of at least 3 biological replicates.

### CK1α-SACK1G co-degradation mediated by DEG-77 requires a direct interaction between CK1α and SACK1G

We have so far shown that degradation of SACK1 proteins by lenalidomide-derived degraders requires CK1α to be present in cells. To definitively demonstrate that it is the interaction between CK1α and SACK1 proteins that facilitates their co-degradation at the endogenous level, we employed skin fibroblast cells derived from a palmoplantar keratoderma (PPK) patient harbouring the homozygous SACK1G^R265P^ mutation that does not interact with CK1α ^14^. Skin fibroblasts derived from an age and sex-matched patient with no PPK pathogenesis served as a wild type control. Loss of CK1α interaction and subsequent inhibition of WNT signalling underlies the pathogenesis of PPK caused by this SACK1G^R265P^ mutation (**Fig. 6A**) ^14^. Fibroblasts from a PPK patient (FIB04), or those from a wild type control (FIB03) were treated with potent CK1α degraders DEG-35 and DEG-77, as well as a weaker degrader DEG-48, for 24 h (**Fig 6B**). As seen with other cell types before, both DEG-35 and DEG-77 caused a robust degradation of CK1α in PPK patient FIB04 as well as FIB03 control fibroblasts, while DEG-48 treatment did not cause any degradation of CK1α in any cells (**Fig. 6B**). In FIB03 control fibroblasts, both DEG-35 and DEG-77 caused a robust depletion in levels of SACK1G compared to DMSO control (**Fig. 6B**). However, despite DEG-35 and DEG-77 causing efficient degradation of CK1α in PPK FIB04 fibroblasts, there was no measurable co-depletion of SACK1G^R265P^ (**Fig. 6B**), indicating that the loss of CK1α binding protects SACK1G^R265P^ mutant from co-degradation by imides. In contrast, in both PPK patient FIB04 and control fibroblast FIB03 cells, DEG-35 and DEG-77 caused a reduction in levels of hyperphosphorylated SACK1D (**Fig. 6B**), suggesting that the CK1α-interacting SACK1 proteins in these cells are potentially still susceptible to co-degradation. The expression of SACK1F and SACK1B was not detectable in these fibroblast cells. To consolidate these findings further, we restored the stable expression of SACK1G-GFP or SACK1G^R265P^-GFP in SACK1G^-/-^ DLD1 cells ^10^. Treatment of these cells with DEG-77 caused a robust reduction in levels of CK1α and SACK1F (**Fig. 6C**). In contrast, while DEG-77 caused a robust reduction in levels of SACK1G-GFP restored in SACK1G^-/-^ DLD1 cells, it failed to deplete SACK1G^R265P^-GFP (**Fig. 6C**). Collectively, these results suggest that SACK1 proteins are only co-degraded when recruited to CUL4A^CRBN^ E3 ligases via their association with CK1α.

**Figure 6:**
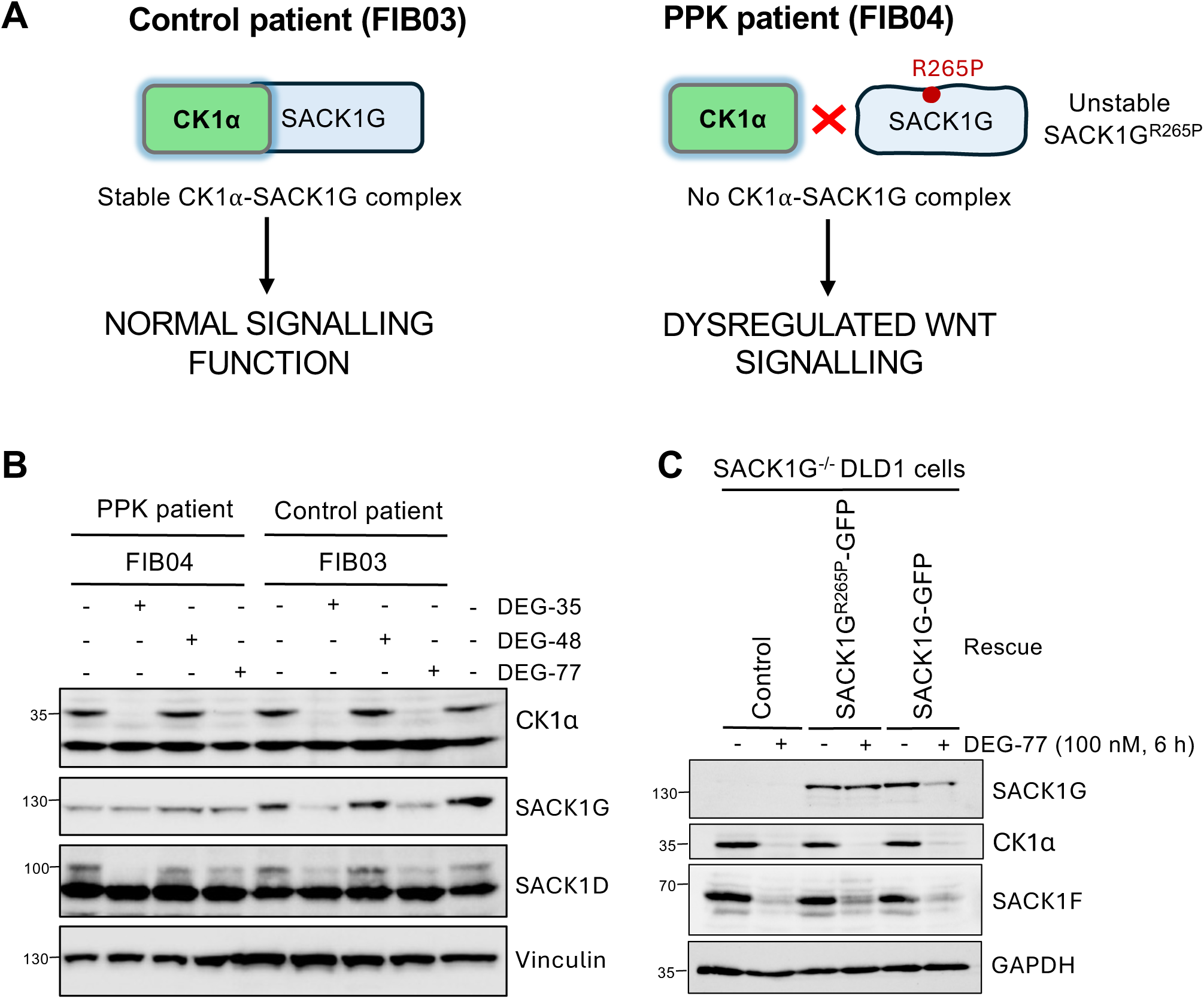
CK1α-SACK1(A-H) co-degradation mediated by DEG-77 and SJ3149 requires a direct interaction between CK1α and SACK1(A-H) proteins. (A) Depiction of CK1α and SACK1G interaction that drives WNT signalling in normal cells (as in control fibroblasts, FIB03) and how this becomes disrupted with the pathogenic SACK1G^R265P^ mutation identified in a patient with palmoplantar keratoderma (as in PPK patient fibroblasts, FIB04). (B) Patient-derived fibroblast cells (FIB04 represents the PPK patient cells harbouring the pathogenic SACK1G^R265P^ variant while FIB03 represents an age and sex-matched control cell line) were treated with 100 nM of DEG-35, DEG-48, DEG-77 or DMSO for 24 h before cell lysis. 10 µg extract protein was resolved by SDS-PAGE and transferred to PVDF membrane for immunoblotting analysis. (C) SACK1G^-/-^ DLD1 cells and those retrovirally transduced to stably express either SACK1G-GFP or SACK1G^R265P^-GFP were treated with DEG-77 (100 nM) or DMSO for 6 h before lysis. 20 µg extract protein was resolved by SDS-PAGE and transferred to PVDF membrane for immunoblotting analysis. For (B&C), blots are representative of at least 3 biological replicates.

## Discussion

Lenalidomide-derived DEG-77 ^37^ and SJ3149 ^40^ are highly potent CK1α degraders. CK1α in cells is localized to unique subcellular compartments via its interaction with one of the eight SACK1 proteins, SACK1(A-H) ^11,13^. Here, we report that DEG-77 and SJ3149 not only degrade CK1α in multiple cell lines but also efficiently co-deplete SACK1F, SACK1G and mitotic SACK1D, with similar potency and kinetics, while some co-depletion of SACK1B was also observed. In contrast, lenalidomide induced poor degradation of CK1α, SACK1B, SACK1D, SACK1G and SACK1H but degraded SACK1F completely, confirming the robust degradation of only CK1α-SACK1F complex ^31^. The depletion of CK1α and the associated SACK1 proteins caused by DEG-77 and SJ3149 was prevented in CRBN^-/-^ cells and by the proteasome inhibitor MG132, suggesting CUL4A^CRBN^ E3 ligase-mediated polyubiquitylation and subsequent degradation by the proteasome. Critically, SACK1 protein depletion by DEG-77 and SJ3149 was completely abolished in CSNK1A1^-/-^ cells, confirming CK1α’s essential role in the co-destabilization of its interacting partners. The requirement for CK1α-SACK1G interaction for DEG-77-induced co-degradation was further validated in cells derived from palmoplantar keratoderma patients harbouring the CK1α-binding deficient SACK1G^R265P^ mutation, where DEG-77 completely failed to degrade SACK1G^R265P^ while still degrading CK1α.

While definitive proof that DEG-77 and SJ3149 induce co-degradation of CK1α-SACK1A-H complexes requires demonstrating direct ubiquitylation of CK1α and associated SACK1 proteins by the CUL4A^CRBN^ E3 ligase machinery, substantial evidence supports co-degradation, particularly for SACK1F. Despite genetic ablation of CK1α stabilizing SACK1F in DLD1 cells, all imide compounds that degrade CK1α to varying levels consistently deplete SACK1F levels. This suggests that the membrane localized CK1α-SACK1F complex forms a productive ternary complex with CUL4A^CRBN^ E3 ligase upon imide treatment, leading to degradation rather that stabilization. The differential targeting of SACK1 proteins may explain the varied potency of imides against CK1α degradation. For instance, lenalidomide’s weaker potency in degrading CK1α could be explained by failure of certain CK1α-SACK1 complexes to form productive ternary complex with CUL4A^CRBN^. Indeed, SACK1G, though not degraded by lenalidomide, was identified as the most abundant interactor of CRBN when cells were exposed to lenalidomide ^25^. Further supporting this notion, overexpression of SACK1G in cells attenuates the ability of lenalidomide to degrade CK1α-SACK1F complexes ^31^.

The co-degradation of CK1α-SACK1G and CK1α-SACK1D complexes by MGDs requires careful interpretation, unlike that of SACK1F. The stability of CK1α and SACK1G proteins is co-dependent. Therefore, degradation of either would lead to the destabilization of both. The mechanisms underlying this co-stability remain unresolved. Interpreting SACK1D co-degradation is also complicated. SACK1D transcription and protein abundance is enhanced in mitosis, where it exclusively interacts with CK1α. This interaction results in SACK1D hyperphosphorylation, which is observed as an upward 25 kDa mobility shift on immunoblots ^6^. Although the mitotic hyperphosphorylated SACK1D disappears upon DEG-77 treatment, this could result from one of three possibilities: i) degradation of CK1α prior to mitosis leaves no CK1α for mitotic SACK1D phosphorylation; ii) the degraders inhibit cells from entering mitosis, which prevents SACK1D from binding CK1α; and iii) CK1α-SACK1D complexes form in mitosis and are then co-degraded. To definitively investigate co-degradation, future experiments should involve treating cells arrested in mitosis with DEG-77, provided CUL4A-CRBN E3 ligase activity is maintained under these conditions.

The glutarimide ring of the IMiD defines IMiD-CRBN interactions whereas the exposed phthalimide group mediates binding to the neo-substrates, dictating the nature of the resulting ternary complex ^47^. Rational design of the DEG series, involving differential additions to the phthalimide amine, yielded compounds with a range of CK1α degradation capabilities ^38^ **(Fig. S1)**. Intriguingly, SJ3149, which achieved similar or better degradation of CK1α compared to DEG-77, degraded SACK1 proteins less efficiently. SJ7095, SJ0040 and SJ3149 were also derived from lenalidomide by additions to the amine group of the phthalimide moiety **(Fig. S1)**. However, the structural differences between SJ3149 and DEG-77 may influence the recruitment of specific CK1α-SACK1(A-H) complexes to CRBN and therefore produce slight structural differences in the resulting ternary complex, potentially accounting for the observed differences in the co-degradation of the SACK1 proteins. While structures of some compounds with CRBN and CK1α have been reported ^35,39,47^, there are currently no structures that also include CK1α with the interacting SACK1A-H proteins.

Because many proteins in cells exist in macromolecular complexes, often defined by post-translational modifications, how degraders recruit the POI macromolecular complex and present the POI or the interacting partner surface to the E3 ligase determines whether the POI and/or the interactor are ubiquitylated and degraded. Here, we provide evidence of *‘bystander degradation’*, where both CK1α and the interacting SACK1B, SACK1D, SACK1F and SACK1G proteins are robustly degraded by DEG-77 and SJ3149. Selective targeting of CK1α catalytic activity has been a major challenge in therapeutics against CK1α-driven malignancies, in part because of the pleiotropic roles of CK1α and the similarity of its kinase domain with other CK1 family members ^5^. SACK1(A-H) provide a subcellular and functional context to CK1 isoforms and, therefore, provide opportunities by which specific CK1 functions can be selectively targeted. Although the DEG and SJ series of compounds assessed here did not display any selectivity towards specific CK1α-SACK1(A-H) complexes, they show that complexes other than CK1α-SACK1F can be targeted using MGDs and, in addition to lenalidomide, selective degraders of these other complexes may be achievable. An alternative approach using PROTACs directed towards individual SACK1 proteins may also be explored as a viable strategy of selectively targeting CK1α function without affecting its other pleiotropic roles.

Our work demonstrating differential CK1α-SACK1(A-H) degradation by different MGDs highlights the importance of protein and cellular context to TPD. SACK1(A-H) are considered the scaffold-like molecular bridges that deliver CK1 isoforms to their substrates. This type of regulation is inherent to many multi-protein assemblies that regulate almost all cellular processes. Given the proximity-inducing nature of small molecule degraders and the role of protein-protein interactions in TPD evidenced here, the underappreciated aspect of bystander degradation, where neighbouring proteins that form a complex with the POI are also targeted for degradation, provides both challenges and opportunities in developing targeted protein degraders.

## Acknowledgements

We thank Prof. Zoran Rankovic (ICR; previously St. Jude) for generously providing us with the SJ series of compounds used in this study. We thank Amaia Lasa-Aranzasti (Vall d’Hebron) for sharing with us PPK and control patient skin fibroblasts used in this study. This study was funded by the UKRI Medical Research Council (MRC), awarded to GPS (MC_UU_00018/6 and MC_UU_00038/6). GPS is also supported by Boehringer-Ingelheim through the Division of Signal Transduction Therapy. CMW and NC are supported by funding from the Department of Defense U.S. Army Medical Research Acquisition Activity (HT94252310263), NIH NCI Spore (5P50CA240243), and the Starr Foundation. LG and BLC are funded by the MRC PPU PhD studentship. TC and KD were supported by UKRI MRC funds awarded to GPS. We thank members of the GPS and CMW labs for helpful discussions. We thank E. Allen, J. Stark and A. Muir for assistance with tissue culture, the staff at the DNA Sequencing services (School of Life Sciences, University of Dundee) and the cloning, antibody and protein production teams within the MRC PPU Reagents & Services (University of Dundee) coordinated by J. Hastie.

## Author Contributions

LG performed most of the experiments, collected and analysed data and contributed to the writing of the manuscript. NC, BLC, & TC performed some experiments and analysed data. KD generated CSNK1^-/-^ and CRBN^-/-^ DLD1 cells. NTW generated some constructs used in this study. TJM designed the strategies, and generated constructs, for CRISPR/Cas9 genome editing experiments. NC and CMW generated the DEG-series of compounds used in this study, analysed data and contributed to the writing of the manuscript. GPS conceived the project, analysed data and contributed to the writing of the manuscript. All co-authors reviewed the manuscript.

## Lead Contact

Further information and reasonable requests for resources and reagents should be directed to and will be fulfilled by the Lead Contact, Gopal Sapkota.

## Data and Code Availability

The original data generated during this study are publicly available at Mendeley Data (link will be added at the time of publication).

## Materials and Methods

### Materials

#### Cell Lines

| Cell Type | Species | Source |
| --- | --- | --- |
| Wild-type DLD1 | Human | ATCC (CCL-221) |
| Wild-type U2OS | Human | ATCC (HTB-96) |
| Wild-type A549 | Human | ATCC (CCL-185) |
| Wild-type TOV-21G | Human | ATCC (CRL-3577) |
| CRBN<sup>-/-</sup> DLD1 | Human | 31 |
| CSNK1A1<sup>-/-</sup> DLD1 | Human | This paper |
| SACK1G<sup>-/-</sup> DLD1 | Human | 31 |
| FIB03 | Human | 14 |
| FIB04 | Human | 14 |

#### Antibodies

| Antibody | Use | Source | Cat No. | Species | Conditions |
| --- | --- | --- | --- | --- | --- |
| CK1α | IB / IP | MRC PPU<br>Reagents and<br>Services | SA527 | Sheep | 1:1000 in 5%<br>milk + TBS-T<br>(IB) 1 μg/mg<br>protein (IP) |
| SACK1B | IB | MRC PPU<br>Reagents and<br>Services | SA270 | Sheep | 1:1000 in 5%<br>milk + TBS-T |
| SACK1D | IB | MRC PPU<br>Reagents and<br>Services | SA102 | Sheep | 1:1000 in 5%<br>milk + TBS-T |
| SACK1F | IB | MRC PPU<br>Reagents and<br>Services | SA103 | Sheep | 1:1000 in 5%<br>milk + TBS-T |
| SACK1G | IB | Abcam | Ab121750 | Rabbit | 1:1000 in 5%<br>BSA + TBS-T |
| SACK1H | IB | MRC PPU<br>Reagents and<br>Services | SA273 | Sheep | 1:1000 in 5%<br>milk + TBS-T |
| CRBN | IB | CST | 71810 | Rabbit | 1:1000 in 5%<br>BSA + TBS-T |
| GAPDH | IB | Proteintech | HRP-60004 | Mouse | 1:5000 in 5%<br>milk + TBS-T |
| Vinculin | IB | Abcam | Ab129002 | Rabbit | 1:1000 in 5%<br>BSA + TBS-T |
| IgG control | IP | MRC PPU<br>Reagents and<br>Services | SPI166 | Sheep | 1 μg/mg<br>protein |

#### Plasmids

| Protein Expressed | Vector | Source | Identifier |
| --- | --- | --- | --- |
| SACK1G-GFP | pBabeD | MRC PPU<br>Reagents and<br>Services | DU29088 |
| SACK1G<sup>R265P</sup>-GFP | pBabeD | MRC PPU<br>Reagents and<br>Services | DU71791 |
| Empty vector | pBabeD | MRC PPU<br>Reagents and<br>Services | DU33769 |

#### Compounds

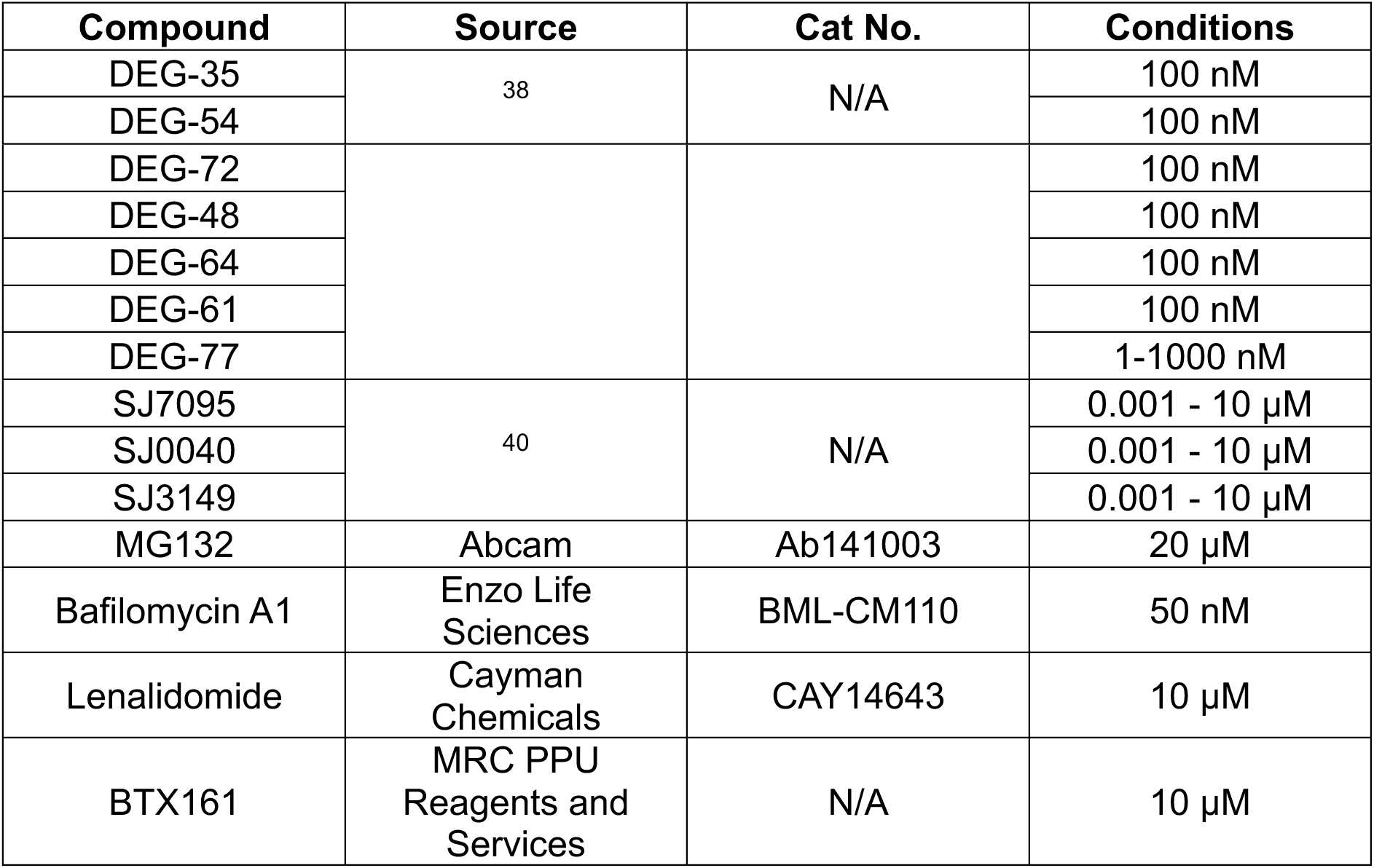

## Methods

### Mammalian Cell Culture

Cells were cultured in either Dulbecco’s modified Eagle Medium (DMEM) (DLD1, U2OS, A549) or Rosewell Park Memorial Institute (RPMI) 1640 medium (TOV-21G) supplemented with 10% (v/v) foetal bovine serum (FBS) except FIB04 and FIB03 fibroblasts which were maintained in DMEM + 20% FBS. Media were also supplemented with 2 mM L-glutamine (Lonza), 100 U/mL penicillin (Lonza) and 0.1 mg/mL streptomycin (Lonza) before use with all cell types. Cells were stored in a humidified incubator at 37°C with 5% CO_2_ and handled under aseptic conditions within a laminar flow hood. For passaging, cells were washed in PBS and incubated with trypsin/EDTA at 37°C until detached and transferred to fresh plates.

### Generation of *CSNK1A1*^-/-^ DLD1 cells by CRISPR/Cas9

All CRISPR/Cas9 technology procedures were performed using a dual guide nickase approach as described previously ^48–53^. CK1α knockout DLD1 cells were generated by targeting the *CSNK1A1* locus with sense guide RNA (pBabeD-puro vector, DU57522); GTCCAAGGCTGAATTCATTGT and antisense guide RNA (pX335-Cas9-D10A vector, DU57527); GATCCTGAGAGACGAAGCTGG. For transfection, 1 μg of each guide RNA was diluted in 1 ml OptiMeM (Gibco) and 20 μl polyethylenimine (PEI: 1 mg/ml). The transfection mixture was vortexed for 15 s and incubated for 20 min at room temperature before adding dropwise to a 10-cm diameter dish containing 70% confluent DLD1 cells in complete media. Media was replaced 24 h post transfection with media containing 2 μg/ml puromycin and incubated for 48 h. Surviving cells were cultured in fresh complete medium until single cells were sorted and plated in individual wells of a 96-well plate which was pre-coated with 1% (w/v) gelatin (Sigma) and contained complete conditioned media supplemented with 20% FCS. Viable clones were expanded and screened by immunoblotting for complete knock-out. Only one CK1α knockout clone was identified, and this was then verified by DNA sequencing. Genomic DNA was isolated using DNeasy Blood & Tissue kit (69505, Qiagen). Primers were generated to amplify the region surrounding the guide RNA target sites with the following primer pairs: CSNK1A1 exon 1 (Forward: AGTAACAGGTAACACCTGCTGAGC; Reverse: AGACATTAGCAAAACTCCAAGTCGC) The region was amplified by polymerase chain reaction (PCR) using KOD Hot Start Polymerase (Merck) according to manufacturer’s instructions. The PCR products were visualised on a 1.5% agarose gel using SYBR Safe DNA gel stain (Invitrogen) and 1 kbp DNA ladders (Promega). The PCR products of positive clones were then cloned into competent cells using the CloneJET PCR Cloning Kit (Thermo Fisher Scientific) according to the manufacturer’s protocol. Isolation of plasmid DNA and sequencing was performed by the MRC-PPU DNA sequencing and services (http://mrcppureagents.dundee.ac.uk).

### Treatment of cells with compounds

The following compounds in DMSO were added directly to culture medium at the indicated concentrations and incubated with cells for the specified times before cell lysis: DEG-compounds (35, 54, 52, 48, 64, 61, 77 - ^38^, St Jude compounds (SJ7095, SJ0040, SJ3149 - ^40^, lenalidomide (Cayman Chemicals), BTX161 (MRC PPU Reagents and Services), MG132 (Abcam) and bafilomycin A1 (Enzo Life Sciences). An equivalent volume of DMSO was added to cells as a treatment control.

### Retroviral transduction

Retroviral vectors were generated by co-transfecting pBabe-puromycin vectors encoding the desired construct (6 µg), pCMV5-gag-pol (3.2 µg) and pCMV5-VSV-G (2.8 µg) (Cell Biolabs) into a 10 cm diameter dish of ∼ 70% confluent HEK293-FT cells as described previously ^14^. In short, plasmids were resuspended gently in 1 mL Opti-MEM medium before being mixed with 24 µL of PEI (1 mg/mL). The transfection mixture was incubated at room temperature for 20 min and then added dropwise to HEK293-FT cells. Next day, culture medium was replenished for 24 h before the retroviral medium was collected and passed through a 0.45 µm sterile syringe filter. Target cells (∼70% confluent) were transduced with the retroviral medium (typically at 1:1 ratio with normal cell culture medium) containing 8 µg/mL polybrene (Sigma-Aldrich) for 24 hr. After 24h, the medium was replaced with fresh culture medium. The medium was again replaced the next day with fresh medium containing 2 µg/mL puromycin for selection of transduced cells. Those cells that survived puromycin selection after all non-transduced control cells died completely by puromycin treatment were taken forward for subsequent experiments.

### Cell Lysis

Cells were harvested by removing culture media, washing twice with ice-cold PBS and then scraping directly into ice-cold NP-40 cell lysis buffer (50 mM Tris-HCl pH 7.5, 0.27 M sucrose, 150 mM NaCl, 1 mM EGTA, 1 mM EDTA, 1 mM sodium orthovanadate, 10 mM sodium b-glycerophosphate, 50 mM sodium fluoride, 5 mM sodium pyrophosphate and 1% NP-40) supplemented with 1x cOmplete protease inhibitor cocktail (Roche) and 1 mM DTT. Cell lysates were transferred to Eppendorf tubes and incubated on ice for 10 min, then clarified by centrifugation at 14000 rpm for 15 min at 4°C. Protein supernatants were transferred to fresh tubes and protein concentration was measured by Bradford assay as previously described ^54^. For SDS-PAGE analysis, samples were prepared to an equal concentration by normalising with NP-40 lysis buffer and 1X LDS sample buffer supplemented with 2.5% β-mercaptoethanol. Samples were boiled at 95°C for 5 min to allow for protein denaturation before loading.

### SDS-PAGE and immunoblotting

Samples prepared in 1xLDS sample buffer containing equal amounts of protein (10-20 µg) were loaded into 10% polyacrylamide gels or 4-12% NuPAGE™ Midi Bis-Tris gels (Invitrogen) and resolved using 120V for 90-120 min. Protein was then transferred to PVDF membrane and successful transfer was verified by Ponceau S staining. Membranes were blocked in 5% (w/v) non-fat milk (Marvel) in TBS-T (50 mM Tris–HCl pH 7.5, 150 mM NaCl, 0.2% Tween 20) and incubated overnight at 4°C with the appropriate primary antibodies. After incubation, primary antibodies were removed, and membranes were washed using TBS-T before undergoing a second incubation with HRP-conjugated secondary antibodies for 1 h at room temperature. Secondary antibodies were prepared in 5% non-fat milk + TBS-T with goat anti-rabbit IgG (7074, CST, 1:2500) or rabbit anti-sheep IgG (31480, Thermo Fisher Scientific, 1:2500). Membranes were again washed after incubation and then exposed to enhanced chemiluminescent (ECL) substrate for 1 min to generate an HRP-linked protein signal. Protein bands were visualised using the ChemiDoc Imaging System (Bio-Rad).

### Immunoprecipitation

Extract containing 2 mg protein were incubated with 2 µg antibody (either against CK1α or IgG control) for 16 h on a rotating platform at 4°C, while a portion of these extracts was retained as input. Protein G Sepharose beads were then equilibrated using ice-cold NP-40 lysis buffer (supplemented with 1 mM DTT) and prepared as a 50% slurry stock. 20 µL of bead slurry was added to each protein extract/antibody mix and both were incubated together on rotating platform for 1 h at 4°C to capture antibody-bound proteins on beads. Beads were then pelleted by centrifugation (1200 rpm, 5 min) and supernatants were stored as flow-through extracts (FT). Beads were washed three times with 0.5 mL ice-cold NP-40 lysis buffer before they were mixed in 50 µL of 1X LDS sample buffer. Immunoprecipitates (IPs) were released from the beads by boiling samples at 95°C for 10 min. Input and IP samples were then resolved by SDS-PAGE, transferred to PVDF membrane and immunoblotted for analysis.

### Quantification and data analysis

Numerical data representing immunoblot quantification performed by Fiji 1.53q (ImageJ) was processed using Microsoft Excel before fold change values were transferred to GraphPad Prism (v.9.4.0) for graph preparation. Quantification graphs are presented as scatter plots with bars and error bars representative of the mean ± standard deviation (s.d.) overlaid with individual datapoints from three biological replicates.

---

**Figure S1:**
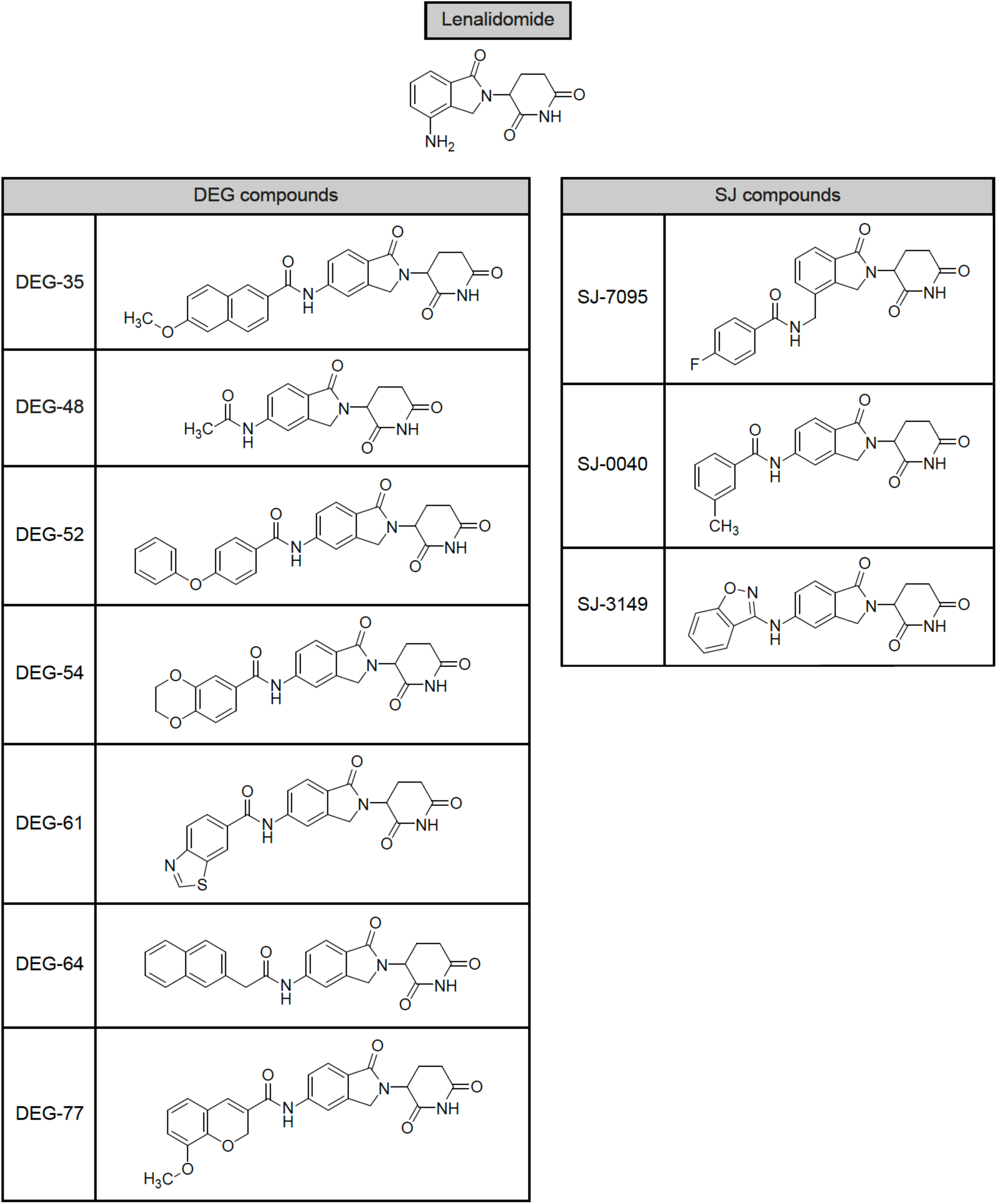
Structures of the DEG- and SJ-series of compounds derived from lenalidomide employed to assess their ability to co-degrade SACK1 domain-containing proteins along with CK1α. The structure of lenalidomide, from which all DEG and SJ compounds derive, is shown at the top. Below this, in the left-hand table, each DEG compound used in this study is listed with its unique chemical structure. The right-hand table depicts the structure of each SJ compound employed in this study.

**Figure S2:**
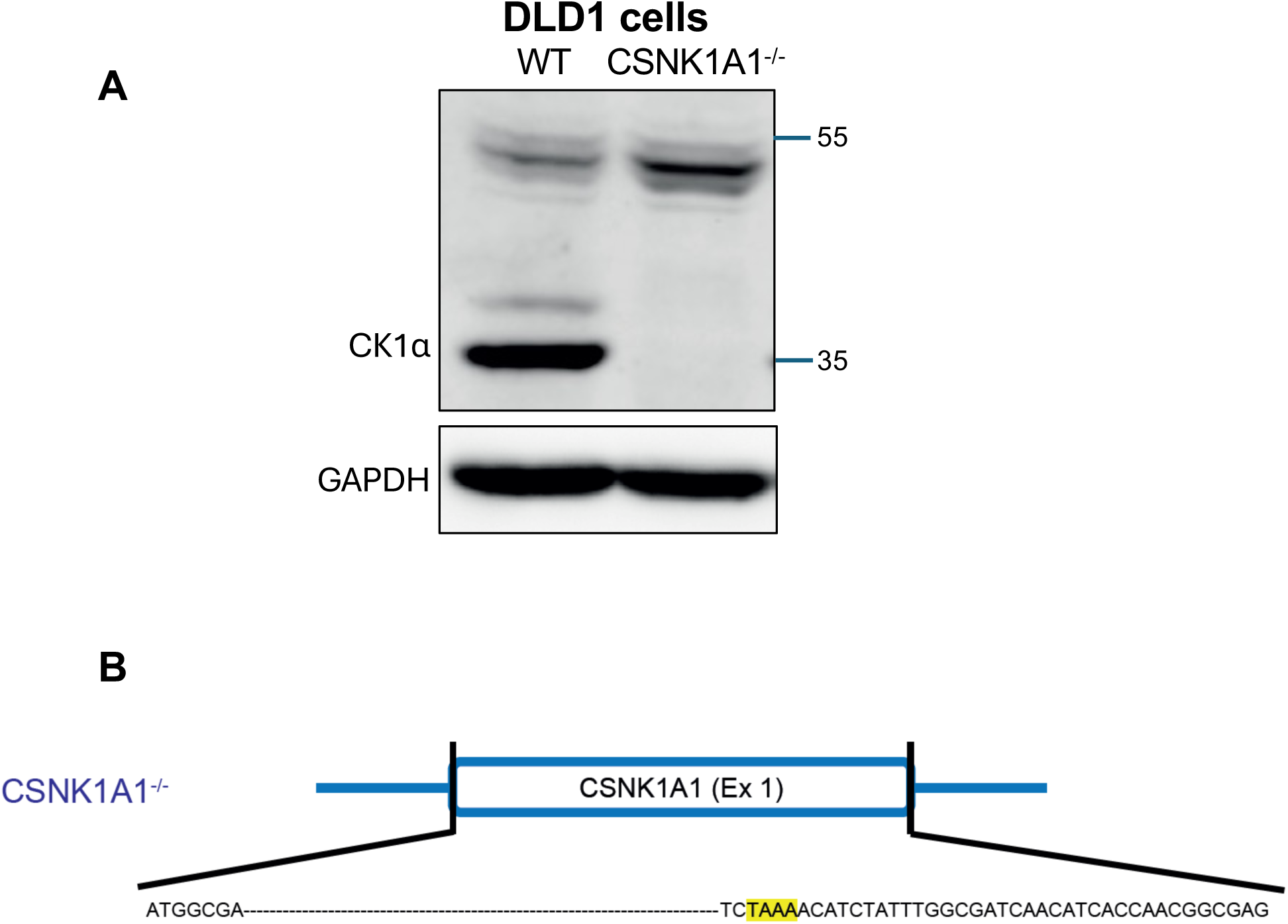
Confirmation of *CSNKA1^-/-^* DLD1 cells. (A) Extracts (20 µg protein) from DLD1 wild-type (WT) and CSNK1A1^-/-^ cells were resolved by SDS-PAGE and subjected to immunoblotting with the indicated antibodies. (B) DNA sequencing of CSNK1A1 (exon 1) confirming alterations in CSNK1A1^-/-^ alleles. Nucleotide deletions and single nucleotide polymorphisms (highlighted nucleotides) predict premature stop codons.

